# A geographical cline in craniofacial morphology across populations of Mesoamerican lake-dwelling fishes

**DOI:** 10.1101/684431

**Authors:** Amanda K. Powers, Carlos A. Garita-Alvarado, Rocío Rodiles-Hernández, Daniel J. Berning, Joshua B. Gross, Claudia Patricia Ornelas-García

## Abstract

The complex geological history and tropical climate of Mesoamerica create a rich source of biodiversity from which we can study evolutionary processes. Here, we discuss highly divergent forms of lake-dwelling fishes distributed across southern Mexico and Central America, originally recognized as members of different genera (*Astyanax* and *Bramocharax*). Recent phylogenetic studies suggest these morphotypes group within the same genus and readily hybridize. Despite genetic similarities, *Bramocharax* morphs exhibit stark differences in cranial shape and dentition. We investigated the evolution of several cranial traits that vary across morphs collected from four lakes in Mexico and Nicaragua and discovered an ecomorphological cline from the northern to southern lakes. Northern populations of sympatric morphs exhibit similar cranial shape and tooth morphology. Southern populations of *Bramocharax*, however, have more maxillary teeth, larger unicuspid teeth, an elongated snout and a streamlined cranium compared to *Astyanax*. The divergence of craniofacial morphology in southern lakes likely evolved in response to environmental pressures. We discuss the ecological differences across the four lake systems in terms of geological history and trophic dynamics. In summary, our study suggests that *Bramocharax* are likely locally-adapted members derived from *Astyanax* lineages, highlighting the complex evolutionary history of the *Astyanax* genus.

## Introduction

The origin and maintenance of biodiversity in nature continues to be one of the most relevant questions in biology. The evolution of a novel adaptive trait corresponds to a phenotypic change originated by either environmental pressures, spontaneous genetic changes, or a combination of evolutionary forces (West-Eberhard 2003). Further, ecological divergence can contribute to morphological evolution and, in extreme cases, promote speciation (Schluter 1996; Barluenga et al. 2006). In particular, cranial structures in fish are useful morphological metrics for understanding trophic specialization processes. For instance, African cichlids illustrate how dramatic craniofacial differences reflect adaptations to diverse feeding niches (Hulsey et al. 2019).

Similarly, the *Astyanax* genus is characterized by a remarkable ability to rapidly adapt to diverse ecological conditions (Miller et al. 2005; Jeffery 2009; Ornelas-García et al. 2018). A well-studied (and extreme) example of morphological change in response to environmental pressure is the *Astyanax mexicanus* cavefish. Thousands of years ago, surface-dwelling forms of *A. mexicanus* invaded caves through river systems, giving rise to a number of modified cave morphs well adapted to the hypogean habitat (Protas et al. 2007; Gross et al. 2009; Jeffery 2009; Elipot et al. 2014; McGaugh et al. 2014). Lacustrine (lake-dwelling) species of the *Astyanax* genus have also evolved morphological changes relative to ecological distribution, including body size and shape changes (Ornelas-García et al. 2008; Ornelas-García et al. 2014; Garita-Alvarado et al. 2018).

Here, we considered four independent lineages of the *Astyanax* genus which present a pair of lake-dwelling morphotypes distributed across southern Mexico and Central America. These two morphotypes are so divergent that they were originally considered different genera (*Astyanax* and *‘Bramocharax’:* Gill and Bransford 1877, Rosen 1970; 1972, Contreras-Balderas and Rivera-Teillery 1983, Bussing 1998). However, recent phylogenetic studies have shown that *Astyanax* and *Bramocharax* indeed group within the same genus (i.e., *Astyanax;* Ornelas-García et al. 2008; Schmitter-Soto 2016; 2017), suggesting these sympatric species are evolving in parallel (Ornelas-García et al. 2008; Garita-Alvarado et al. 2018).

Despite genetic similarity, morphological differences between lake-dwelling *Astyanax* and *Bramocharax* have long been appreciated (Gill and Bransford 1877). Geometric morphometric analyses revealed major differences between the two morphs relating to tooth shape, eye size, snout length, body depth, head profile and mouth orientation (Ornelas-García et al. 2014). *Astyanax* morphs are characterized by a deep body, blunt snout and multicuspid teeth compared to the fusiform (narrow) bodied, long-snouted, unicuspid *Bramocharax* (Ornelas-García et al. 2014). Interestingly, a recent body shape and dentition analysis (Garita-Alvarado et al. 2018) revealed morphological differences between morphs vary across localities (i.e. geographically-distinct lakes). Based on this genetic and morphological data, we hypothesized that members of the genus *Bramocharax* are actually locally-adapted morphs within lineages of *Astyanax*.

In order to test this idea, we assessed morphological divergence in the two morphs sampled from four Mesoamerican lakes with different geological and ecological histories. We evaluated nine variables based on skull morphology and dentition using a combination of traditional and three-dimensional (3D) geometric morphometric analyses. We discovered significant morphological changes that reflect a geographical cline from northern to southern lakes. Further, we discuss the unique geological history and trophic dynamics of each lake system and how this plays a role in local adaptation of *Astyanax* species. Taken together, our study illustrates the complex evolutionary history of lake-dwelling Mesoamerican characids and clarifies the long-debated characterization of species within the *Astyanax* genus.

## Materials and Methods

### SPECIMEN COLLECTION

Adult, wild-caught fish corresponding to the two morphotypes originally recognized as different genera (i.e. *Astyanax* and *Bramocharax)* were collected from four lakes in Mexico and Nicaragua (Fig. 1), North to South: Lake Catemaco, in the Papaloapan basin Mexico *(Astyanax-like* n= 2, *Bramocharax* n= 3); Lake Ocotalito and the Lacanjá river in the Usumacinta basin, Mexico *(Astyanax* n= 3, *Bramocharax* n= 12); Lake Managua (also known as Xolotlán), in the San Juan basin Nicaragua *(Astyanax* n= 4, *Bramocharax* n= 4) and Lake Nicaragua (also known as Cocibolca or Granada), in the San Juan basin, Nicaragua *(Astyanax* n=4, *Bramocharax* n= 5) (Table S1). We included individuals from four species originally assigned to the genus *Bramocharax: B. caballeroi* from Lake Catemaco (Contreras-Balderas and Rivera-Teillery 1985), *B. bransfordii* from Lake Nicaragua and the Sarapiquí River (Gill and Bransford 1877) Lake Nicaragua, *B. elongatus* from Lake Managua (Meek 1907), and *Astyanax ocotal* (considered Bramocharax-like) from Lake Ocotalito (Valdez-Moreno et al. 2009, Schmitter-Soto 2017). We also evaluated three species assigned to the genus *Astyanax: A. nicaraguensis* from Lake Nicaragua and the Sarapiquí River (Eigenmann and Ogle 1907), *A. nasutus* from Lake Managua (Meek 1907), and *A. aeneus* from Lake Catemaco and Lake Ocotalito (Ornelas-García et al. 2008; Schmitter-Soto, 2017). Sampling was carried out between 1997-2015, using different diameter mesh-size nets. Fish were euthanized in ice-water according to the protocol in the approved SGPA/DGVS/02438/16 collection permit. Fish were collected under permits 007-2013-SINAC and SINAC-GASP-PI-R-072-2014, Sistema Nacional de Áreas de Conservación and Ministerio de Ambiente y Energía of Costa Rica (Costa Rica), 008-112014/DGPN, Ministerio del Ambiente y los Recursos Naturales (Nicaragua), and SGPA/DGVS/05464/, SAGARPA (México). Voucher specimens were preserved in 95% ethanol and deposited in the Colección Nacional Peces, Instituto de Biología, UNAM (CNPE, IBUNAM) and the ichthyological collection of El Colegio de la Frontera Sur (collection numbers: ECOS875, ECOSC-6993, ECOSC-6994, ECOSC-7002, ECOSC-7010, ECOSC-7060). Hereafter the different sympatric morphotypes are referenced by their original genera. For instance, we refer to ‘elongate-body’ morphotype as *Bramocharax* and for the ‘deep-body’ morphotype as *Astyanax*.

**Figure 1.**
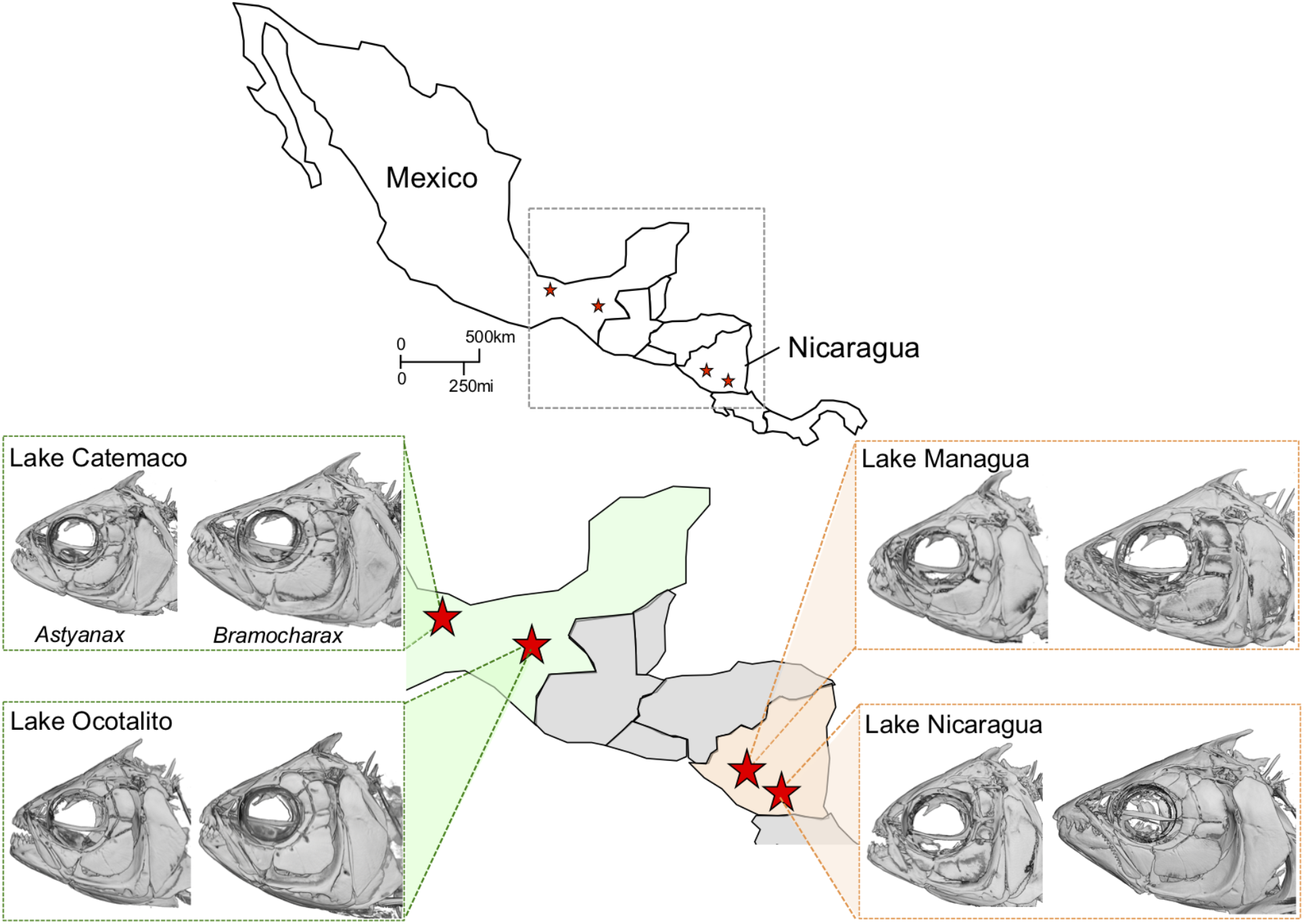
Distribution of *Astyanax* and *Bramocharax* morphs across four Mesoamerican lakes. The *Astyanax* genus comprises of fish species widely distributed across Mexico and Central America. Lake-dwelling morphs demonstrate genetic similarity, but morphological differences in craniofacial and body shape. Here we characterize wild-caught *Astyanax* and *Bramocharax* specimens from four lakes: two Mexican lakes (Catemaco and Ocotalito; green) and two Nicaraguan Lakes (Managua and Nicaragua; orange).

### MICRO-COMPUTED TOMOGRAPHY SCANNING

Specimens were shipped to the University of Cincinnati Vontz Core Imaging Laboratory (VCIL; Cincinnati, Ohio, USA) for micro-computed tomography (microCT) scanning. The Inveon Multimodality System (Siemens; Munich, Germany) was used to collect two-dimensional slices in the axial, coronal and sagittal orientations. Slices were stored as DICOM formatted files and reconstructed into three-dimensional (3D) volume-rendered images (Fig. 1) using Amira software (v6.1; FEI Company, Hillsboro, OR). Bone was labeled based on density using the ‘label field’ tool in Amira and segmented to create a 3D surface file of the skull.

### ECOMORPHOLOGICAL ANALYSES

For the trophic morphology analyses between morphs and between lakes, we measured 36 specimens (Table S1) using the ‘3D length’ measuring tool in Amira: length of the most anterior tooth of right premaxillary (PT), length of the most anterior tooth of right dentary (DT), lateral premaxillary length (WP), dorsal premaxillary length (PP), right opercle length (OL), right opercle width (OW), eye orbit (EL) from lateral ethmoid to superior fourth suborbital bone (SO4) and dorsal foramen width (DF). All measurements were standardized by dividing them by head length. Further, a principal component analysis (PCA) was performed to determine and visualize major axes of variation using the log-transformed variables.

For the analysis of maxillary teeth number, we included a total of 131 specimens from the four lakes. We included 62 newly dissected specimens (Lake Catemaco: n= 1 *Astyanax*, n= 3 *Bramocharax;* Lake Ocotalito: n= 9 *Astyanax*, n= 38 *Bramocharax;* Lake Managua: n= 4 *Bramocharax;* Lake Nicaragua: n= 2 *Astyanax*, n= 5 *Bramocharax)* and 69 specimens from Garita-Alvarado *et al.* 2018 (Table S1). We performed generalized linear modeling (GLM) to compare the number of maxillary teeth (response variable) using a quasi-Poisson distribution in R (R statistical language: package stats). We tested the effect of morph identity, geography, log-SL as covariate and the interaction morph × geographical system. Additionally, a GLM was performed to compare the number of maxillary teeth between morphs in each lake (log-SL and morph as factors, including the Bonferroni correction for several comparisons)

For the analysis of tooth morphology on specimens from Lake Ocotalito, Chiapas, *(Astyanax* n=6, *Bramocharax* n=27), we determined the distribution of the type of the most anterior tooth (i.e. tricuspid or hexacuspid) of the inner row of teeth of the premaxilla and related it with the standard length by morph. To compare the type of the most anterior tooth between morphs, we performed an ordinal logistic regression for each response variable in R (R statistical language: package ordinal). We included type of tooth (tricuspid to hexacuspid) as the ordinal response variable and the predictor variable morph, log SL as covariate and its interaction. Additionally, we compared the distribution of the type of the most anterior tooth of the inner row of teeth of the premaxilla by morphs from Lake Ocotalito with the same distribution by morphs from San Juan and Catemaco systems from Garita-Alvarado et al. (2018).

### ANALYSES OF SKULL SHAPE WITH GEOMETRIC MORPHOMETRICS

Twenty-four homologous landmarks were collected from dorsal, lateral and ventral orientations inclusive of the premaxilla, maxilla, mandible, supraorbital, lateral ethmoid, and opercle bones in 3D for 37 specimens (Table S1). Landmark collection and shape analyses were conducted as described in Powers et al. (2017). Briefly, cranial shape was analyzed using MorphoJ software (Klingenberg 2011) to perform a principal component analysis (PCA) allowing visualization of cranial shape differences across *Astyanax* and *Bramocharax*. Wireframe graphs for principal component 1 (PC1) were created to illustrate changes across the geographical cline. To assess the morphological variation across the morphs and systems, we performed a canonical variate analysis (CVA), and generated wireframe shape changes associated with the canonical variates, scaled by Mahalanobis distance units in MorphoJ software.

## Results

### ECOMORPHOLOGY ACROSS MEXICAN AND NICARAGUAN POPULATIONS

Using the PCA for eight ecomorphological traits, we explored the morphospace across northern (i.e. Catemaco and Ocotalito) and southern (i.e. Managua and Nicaragua) lake populations. The first and second components accounted for 84% of the total variance (56.5% for PC1 and 27.5% for PC2; Fig. 2A). The positive scores of PC1 were related to DF and PP (Fig. 2A; Fig. S1). In PC1 we observed a clear segregation between morphs from Managua and Nicaragua, in contrast to Catemaco and Ocotalito. In the latter, *Astyanax* and *Bramocharax* sympatric morphs completely overlap. In PC2, the positive scores were associated with the DF and PT, while along the negative axis we recovered higher scores associated with WP and PP. The extreme negative PC2 axis is associated with dentary tooth length and lateral premaxillary length of the *Bramocharax* morph from Managua and Nicaragua (Fig. S1). Interestingly, we found a decrease in the size of the premaxillary teeth in *Bramocharax* morphs from lakes Managua and Nicaragua (Fig. S1), but an increase in the size of the lower jaw dentary teeth in morphs from lake Nicaragua (Fig. S1).

**Figure 2.**
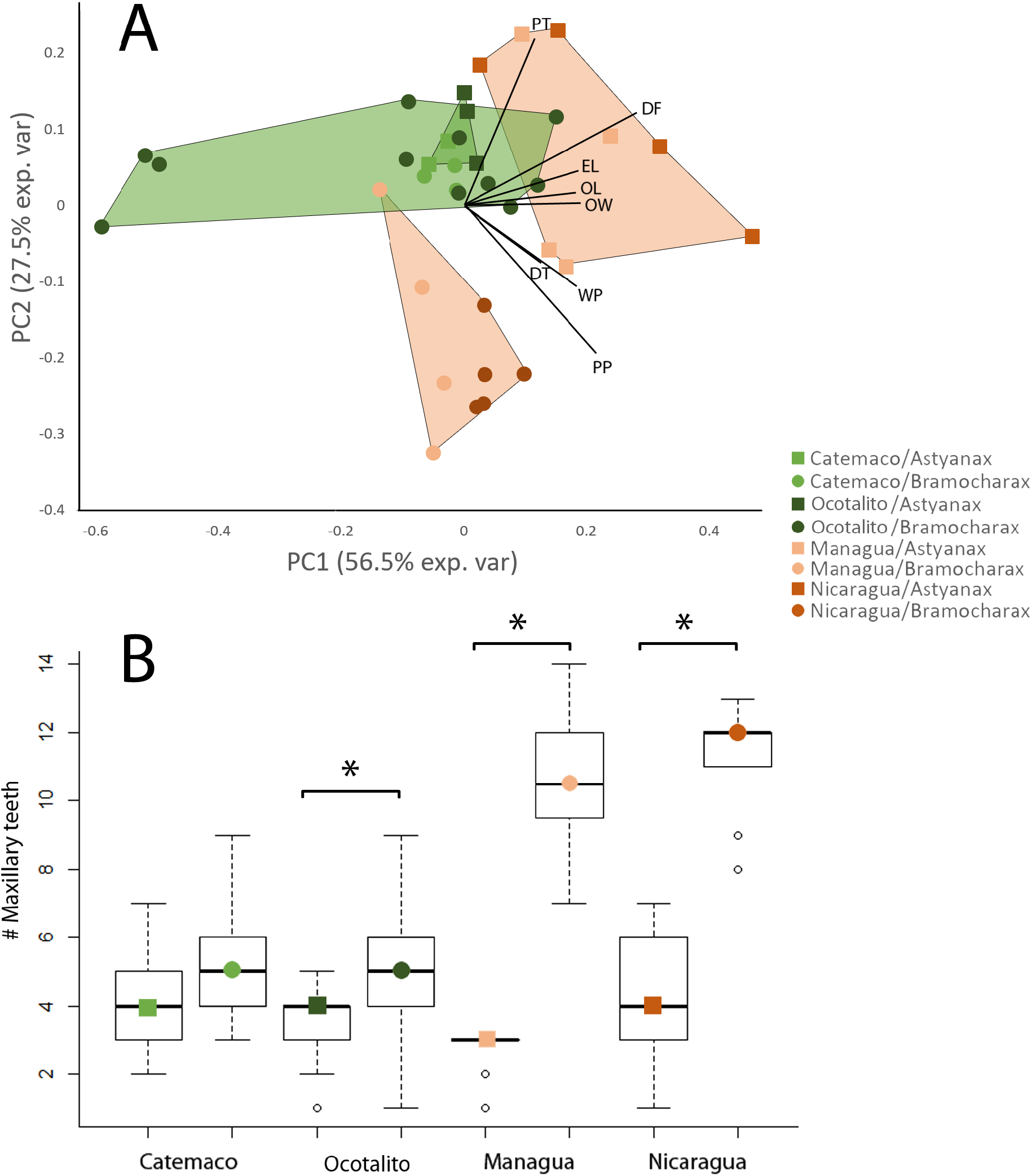
Principal component analysis (A) of trophic-related traits by geography and morph with loadings for principal components 1 and 2 (PT: right premaxillary tooth length, DT: right dentary tooth length, WP: lateral premaxillary length, PP: dorsal premaxillary length, OL: right opercle length, OW: right opercle width, EL: Eye orbit from lateral ethmoid to superior SO4 and DF: dorsal foramen width). Boxplot of number of maxillary teeth by morph and system (B).

For number of maxillary teeth, differences varied depending on the morph and lake combination: *Bramocharax* morph showed more maxillary teeth than *Astyanax* with a higher disparity in Lake Nicaragua and Lake Managua (southern populations) than in Lake Catemaco and Lake Ocotalito (Fig. 2B). Also, the number of maxillary teeth was positively correlated with log SL (morph: F1=114.8, p<0.001, geographic system (i.e. lake): F3=41.4, p<0.001, log SL: F1 =16.1, p<0.001, morph*geographic system: F1=15.8, p<0.001, Fig. S2). Within lakes, we found differences between morphs in all paired comparisons except between Catemaco morphs (Nicaragua: F= 91.6, p<0.001; Managua: F= 190.7, p<0.001; Catemaco: F= 4.12, p= 0.1, Ocotalito: F=9.76, p<0.001).

In contrast to convergence of multicuspid tooth patterning across *Astyanax* populations, we observed divergence in the number of cusps in northern and southern populations of *Bramocharax*. In the southern lake populations, anterior premaxillary teeth were almost always unicupid (canine) for *Bramocharax* morphs (Fig. S2). The Catemaco lake population, however, generally exhibit tricuspid teeth. For specimens from Lake Ocotalito, we found no differences in the type of the most anterior tooth of premaxilla between morphs (z=1.5, p=0.13). Further, the type of the most anterior tooth and log SL were not correlated (log SL: z=1.2, p=0.21 and morph*log SL interaction: z=-1.4, p=0.14) (Fig. S2). In the *Bramocharax* morph, the type of the most anterior tooth of the inner row of teeth of the premaxilla varied from tricuspid to hexacuspid, while the *Astyanax* morph did not show tricuspid teeth. Hexacuspid was the most common type of the most anterior tooth of the inner row of teeth of the premaxilla followed by pentacuspid in the *Bramocharax* morph; the *Astyanax* morph showed the same relative abundance for all types of the most anterior tooth (Fig. S2).

### SKULL SHAPE DIFFERENCES ACROSS MEXICAN AND NICARAGUAN POPULATIONS

Using 3D geometric morphometrics, we examined global skull shape in both morphs across northern (i.e. Catemaco and Ocotalito) and southern (i.e. Managua and Nicaragua) populations. The PCA revealed that northern populations of both *Astyanax* and *Bramocharax* occupy a similar morphospace (PC1 = 41.3% shape variation, PC2 =12.9%), while southern populations have diverged in terms of global cranial shape (Fig. 3A). PC1 illustrates the major differences between the two morphs. *Bramocharax* morphs have a longer head, elongated jaws, a narrower head and an upwardly-projected mouth. In contrast, the *Astyanax* morphs have a shorter snout, wider head and a lower mouth position (Fig. 3A).

**Figure 3.**
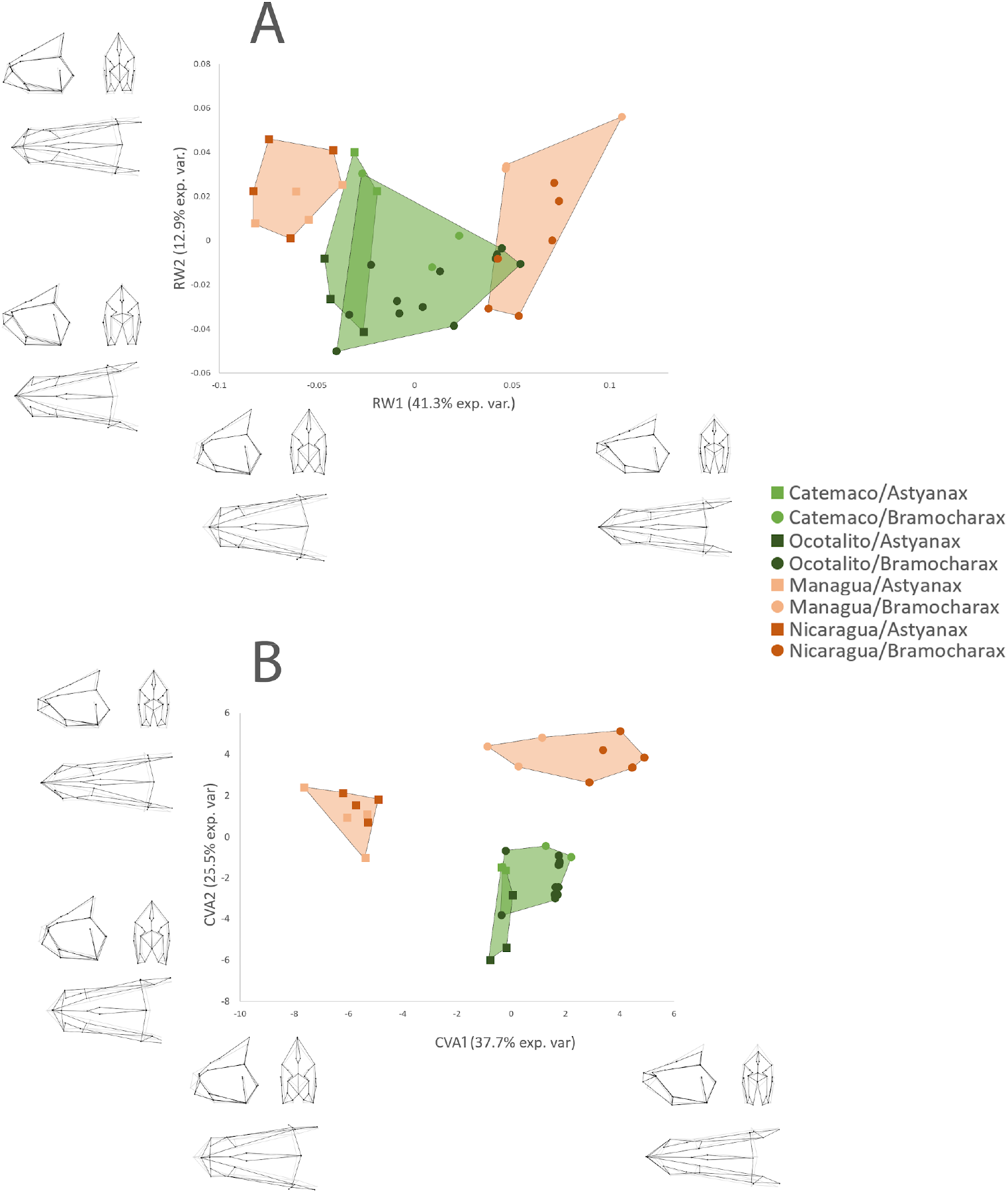
Principal component analysis (A) and canonical variate analysis (B) by lake and morph. Transformations of landmark positions represent the extreme changes of head shape (black lines) along the axes with respect to the consensus shape (gray lines). Hulls denoted north (Lake Catemaco and Lake Ocotalito; green) and south populations (Lake Managua and Lake Nicaragua; orange) by morph.

In the CVA analysis (Fig. 3B), we observed a consistent pattern where the first two canonical variates account for 63.2% of the variance. CV1 differentiates between the two morphs, which demonstrates at the lateral aspect a divergence in head length, snout length and mouth position, and in the ventral axis, we observed a difference in head length, and a difference in skull width was observed in the ventral and frontal axes. CV2 differentiates between the southern and northern systems, however, similar to the CV1, we observed substantial overlap between the two morphs in the northern lake systems (Catemaco and Ocotalito). The southern populations exhibited a dramatic elongation of the upper and lower jaws (premaxilla, maxillary and mandible) compared to northern populations (Fig. 3B).

We observed differences in opercle shape (a downward shift at the posterior point) between northern and southern populations with no significant difference in size (Fig. 4). Eye size, normalized to head length, was consistent across lakes, with larger eyes in *Astyanax* compared to *Bramocharax* morphs (Fig. S1). This ‘streamlined’ cranial shape was also quantified using angle measurements. We found that morphs from lakes Managua and Nicaragua showed a decrease in the dorsal foramen angle and the ventral mandible angle compared to *Bramocharax* morphs from the northern lake Ocotalito population (Table S1). Other shape changes observed in the southern population were posteriorly and medially displaced opercle bone, a decrease in the slope of the dorsal cranium, and larger eyes (Fig. 4).

**Figure 4.**
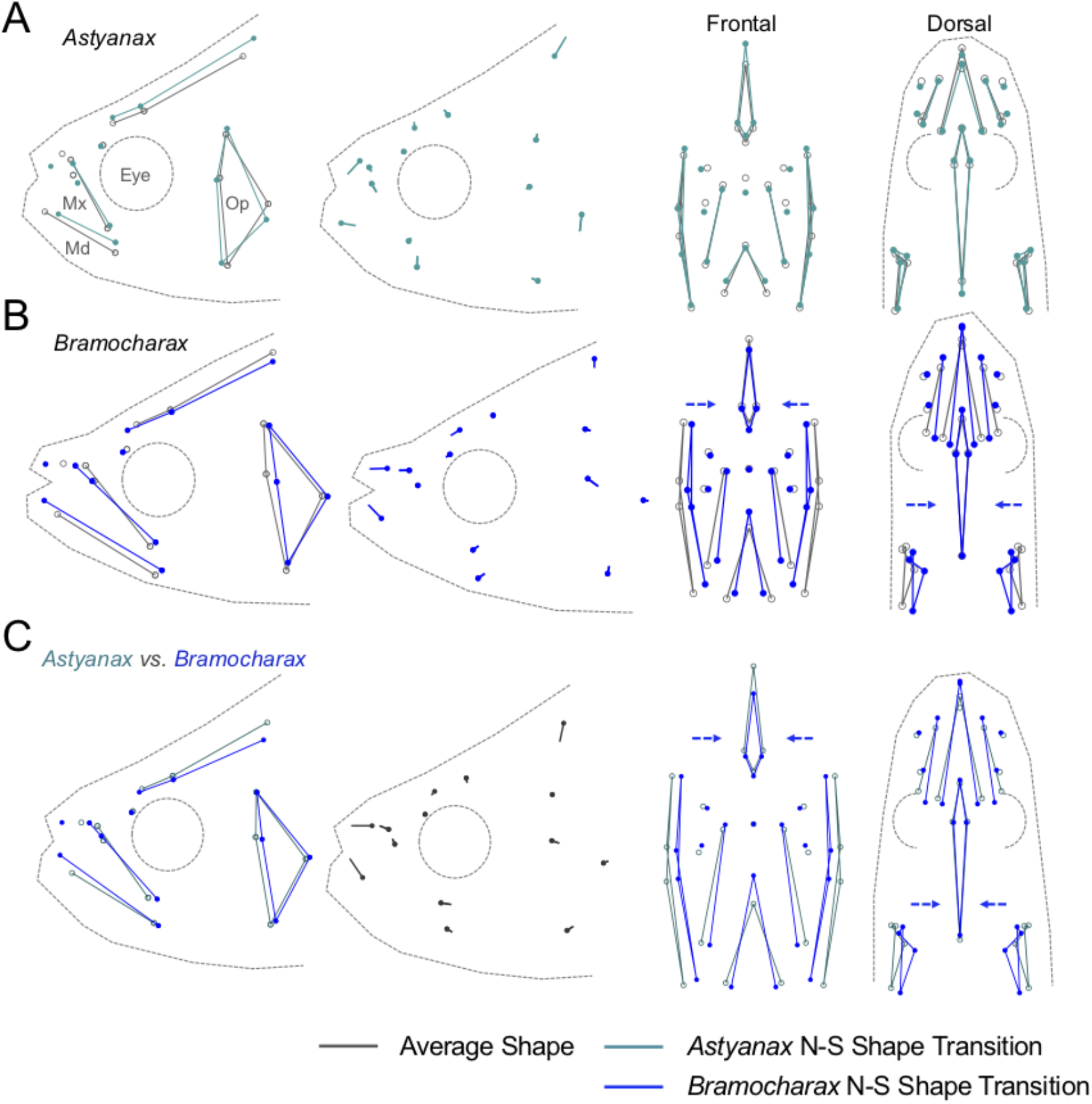
Wireframe graphs for PC1 reveal differences in craniofacial shape across a geographical cline. Southern populations of *Astyanax* exhibit modest shape differences in the opercle (Op) and mandible (Md) bones (A). In contrast, southern populations of *Bramocharax* illustrate more extreme differences in skull shape, including a lengthening of the mandible (Md) and maxillary (Mx) bones and a narrowing of the craniofacial complex (shown in the frontal and dorsal views; blue arrows; B). Average shape shown in gray. *Astyanax* north-south shape transition shown in teal and *Bramocharax* north-south shape transition shown in blue. Overall shape changes between *Astyanax* and *Bramocharax* morphs were similar to the north-south transition, with an elongated snout and fusiform shape in the *Bramocharax* morphs *(Bramocharax* in shown in blue and *Astyanax* in teal).

## Discussion

### DIFFERENCES IN TROPHIC DYNAMICS MAY EXPLAIN CLINAL VARIATION IN SKULL SHAPE AND DENTITION

We found evidence of an ecomorphological cline in sympatric species of fish across a geographical landscape of lakes in Mexico and Nicaragua. Northern and southern populations of *Astyanax* exhibit minor shape changes in the premaxilla with little appreciable difference in dentition. In contrast, southern populations of *Bramocharax* show striking differences in cranial shape, characterized by an elongation of the jaws and narrow, fusiform skulls compared to populations from northern lakes. Furthermore, *Bramocharax* morphs from southern lakes demonstrated dramatic changes in dentition, with increases in maxillary teeth, tooth size, as well as observed diastemas between teeth, and loss of cusps. These results are consistent with whole-body shape differences observed by Garita-Alvarado et al. (2018).

This morphological disparity likely corresponds to trophic differences between the northern and southern populations. According to stable isotope analyses on nitrogen content of the fish gut, in the northern lakes *Astyanax* and *Bramocharax* morphs showed an overlap in their trophic niches (Ornelas-García et al. 2018). Specifically, in Lake Catemaco, the diet of the *Astyanax* morph consisted of 79% vegetation/detritus, 20% fish, and 1% insects, while the *Bramocharax* morph diet consisted of 61% vegetation/detritus, 24% fish, 15% invertebrates (Ornelas-García et al. 2018). This suggests that in the northern lakes, both morphs are omnivorous, with the *Bramocharax* morph eating a larger proportion of invertebrates. Interestingly, gut content analyses from Central American lakes suggests that southern populations of *Bramocharax* are more piscivorous (primarily eating other fish) compared to northern populations. Bussing (1998) noted that *Bramocharax bransfordi* from Costa Rica, living in close proximity to Lakes Managua and Nicaragua, had 51% fish, 49% invertebrates in the gut with no detectable traces of vegetation during post-mortem dissections.

The similar trophic niches of *Astyanax* and *Bramocharax* morphs in the northern populations is congruent with the higher similarity in their cranial shape and dentition phenotypes. In northern lake populations, even when the *Bramocharax* morph exhibit a fusiform body, larger head and unicuspid teeth relative to the southern *Bramocharax* morphs (Ornelas-García et al. 2014), they have shorter snouts (Fig. 3; Fig. 4A) and more overlap in cranial shape between sympatric morphs (Fig. 2). This is consistent with morphological and trophic dynamics in omnivorous lake-dwelling African cichlids (Streelman and Albertson, 2006). In contrast, the divergence in cranial morphology in the southern populations of both morphs may be attributed to differences in diet, although gut content analysis of southern *Astyanax* populations has yet to be performed.

*Astyanax* morphs in southern lakes resemble their northern counterparts with no significant difference in maxillary teeth number (Fig. 2) and minor shape differences in the jaw with an elongation of the premaxilla and shortening of the mandible (Fig. 4B). In contrast, southern populations of *Bramocharax* exhibit traits consistent with predatory habits, including large, unicuspid teeth, elongated jaws for prey capture (Liem 1978; Cooper et al. 2011; Burress et al. 2016; Bonato et al. 2017) and fusiform (streamlined) shape for accelerated swimming (Porter and Motta 2004; Arbour and López-Fernández 2014; Ornelas-García et al. 2014; Ornelas-García et al. 2018). Similar attributes have been reported in a predator Neotropical characid genus *Oligosarcus* (Santos et al. 2011; Bonato et al. 2017). We also found evidence of change in shape (toward a longer, narrower bone) for the opercle in the southern populations of *Astyanax* and *Bramocharax*. The opercle bone and the opercular complex are important for respiration and feeding (Kimmel et al. 2008), therefore this shape change over the geographical gradient may reflect differences in feeding strategies.

### GEOLOGICAL HISTORY OF MESOAMERICAN LAKE SYSTEMS LIKELY INFLUENCES MORPHOLOGY

The ecomorphological cline in cranial shape and dentition we describe may also be a consequence of the complex geological history in Mesoamerica. Characiforms (the order inclusive of both *Astyanax* and *Bramocharax* morphs) originated in South America (Orti et al. 1997; Otero and Gayet 2001; Calcagnotto et al. 2005; Ornelas-García et al. 2008) and likely dispersed throughout Central America and Mexico through colonization events prior to the late Cenozoic closure of the Panama Strait (3.3 Mya). In this regard, the degree of morphological differences between *Astyanax* and *Bramocharax* morphs within each lake is likely associated with the age of the lakes. The independent origin of the *Bramocharax* morph could be related to both phylogenetic history of the genus, and the degree of divergence could be associated to the geological history of the lake systems (Ornelas-García et al. 2008; Díaz et al. 2017; Stoppa et al. 2018). This would suggest that the Central American populations are the oldest lineages, Lake Managua and Lake Nicaragua were present since the early Pleistocene (~ 2.5-1 Mya; Stoppa et al. 2018), and the Usumacinta region (Lake Ocotalito) corresponds to the youngest lake (at least from ~9,000 years BP, Díaz et al. 2017).

Our results demonstrate Central American populations have the largest differentiation between sympatric morphs. In Mexico, however, even when cranial shape differences and trophic characters diverge in the same direction as the Central American morphs (elongation of the jaws and narrow, fusiform skulls, and an increase in maxillary teeth, tooth size), the disparity between morphs was lower. Similarly, this pattern of disparity dependent on lake age may be linked to feeding morphology divergence in adaptive radiations of African cichlids (Cooper et al. 2010). Our results provide insight to the relationship between the age of the lake systems and the morphological disparity between sympatric morphs in Mexico and Central American lakes. Additionally, this study system provides the unique opportunity to explore the relative roles of adaptive divergence and geographical differences in the skull morphological variation depending on the lacustrine system.

## Conclusion

Contrary to prior historical accounts of morphological divergence between *Bramocharax* and *Astyanax*, we observed similarities between the morphs in two lake populations in Mexico. Further, we discovered an ecomorphological cline across northern and southern populations, with significant morphological divergence in two Nicaraguan lakes. This supports the notion of repeated local adaptations of *Bramocharax* morphs originating from *Astyanax* lineages. This study system provides a unique opportunity to explore convergent patterns in ecomorphological traits associated with repeated evolution. In sum, the morphological differentiation between morphs likely has functional relevance influenced by the complex geological history and trophic dynamics of the lake ecosystems.

## Supporting Information

**Supplemental Table 1.** Morphometrics landmarks and measurements.

A. Locality, Landmark data and species information. B. Geometric morphometrics data, Relative Warps values and Canonical Variates values. C. Meristic data, Morphological measurements, morphological ratios and Log Transformed variables used in the PCA. D. Maxillary teeth number by morph and system. E. Tooth type.

**Supplemental Figure 1 (Fig. S1).**
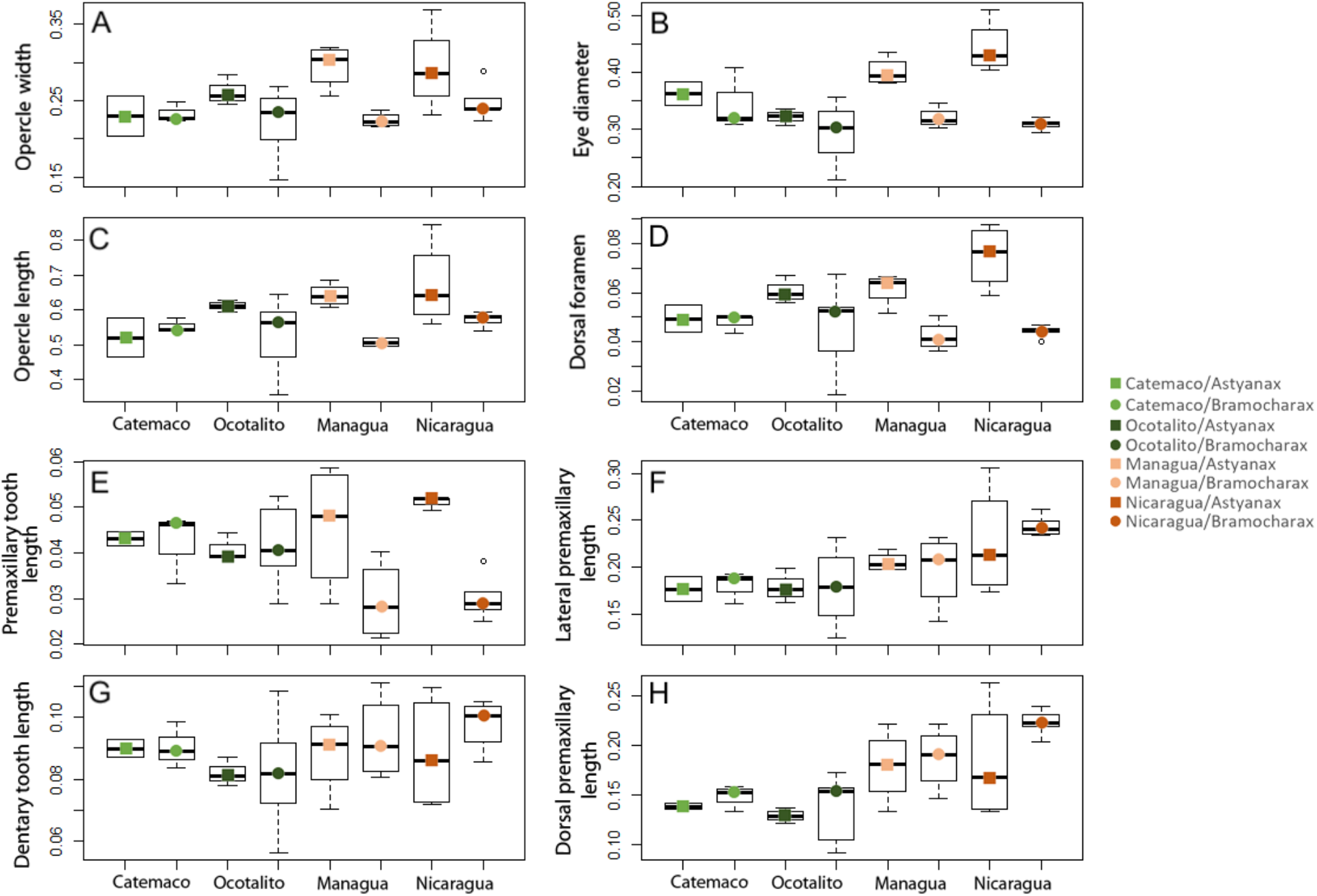
Craniofacial and tooth size metrics across lake systems. Size metrics including opercle width (A), eye diameter (B), opercle length (C), dorsal foramen width (D), premaxillary tooth length (E), lateral premaxillary length (F), Dentary tooth length (G), and dorsal premaxillary length (H) were measured for morphs in each of the four lake systems. Lake systems are coded as follows: Catemaco (light green), Ocotalito (dark green), Managua (light orange) and Nicaragua (dark orange). All measurements are in millimeters.

**Supplemental Figure 2 (Fig. S2).**
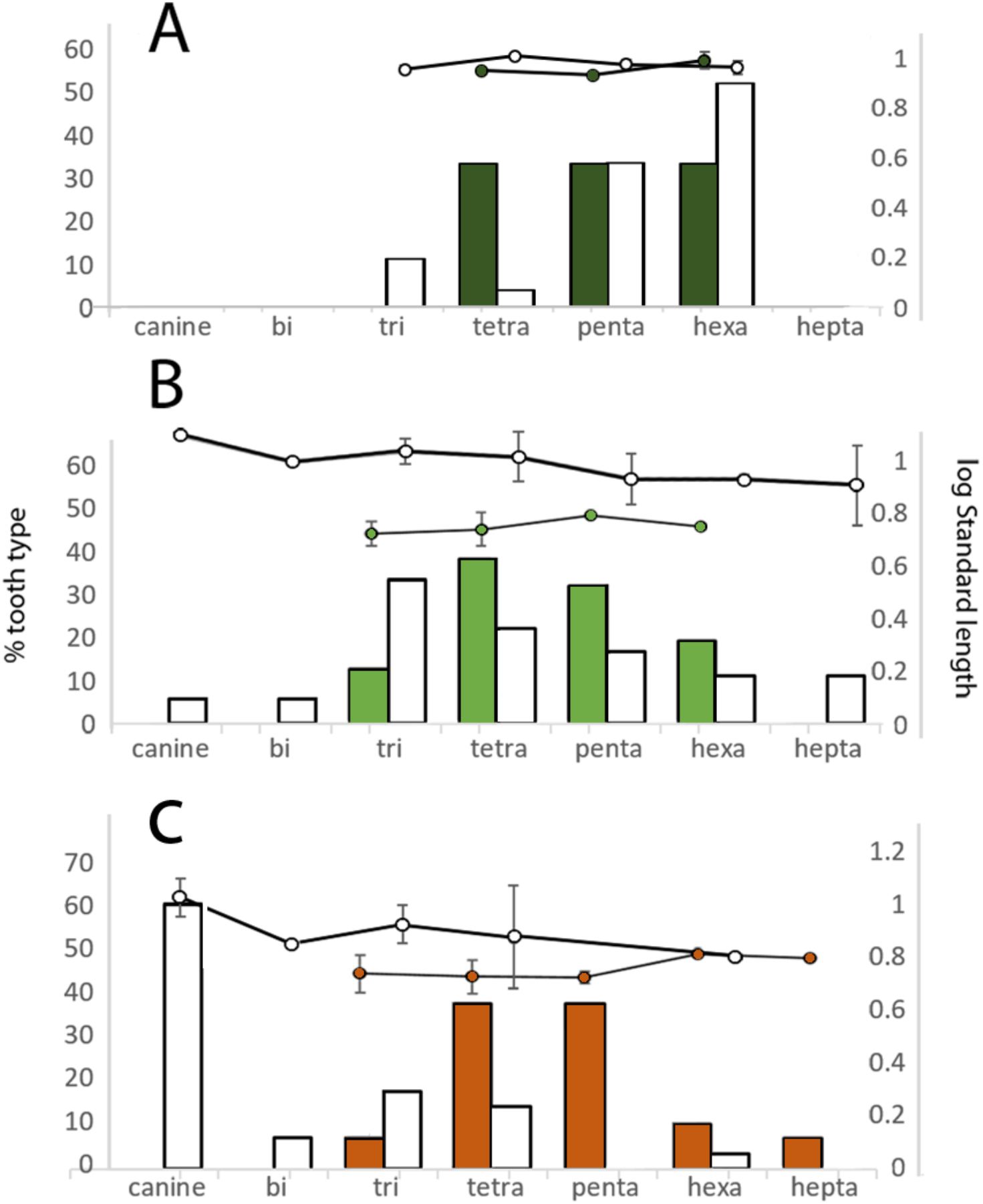
Tooth type frequency across lake systems. Percentage of types of the most anterior tooth of premaxilla (left axes, bars) as defined by tooth cusp number was analyzed for *Astyanax* (colored bars) and *Bramocharax* (white bars) morphs for lakes Ocotalito (A), Catemaco (B) and Nicaragua (C). Log standard length (right axes) illustrated specimen size for *Astyanax* (colored circles) and *Bramocharax* morphs (white circles).

## AUTHOR CONTRIBUTIONS

CPO-G, JBG and AKP conceived the study and designed experiments. AKP, CA-G, DJB and RR-H performed experiments and analyzed data. AKP and CPO-G wrote the manuscript.

## ACKNOWLEDGMENTS

The authors would like to thank members of the Ornelas-García, Gross and Rodiles-Hernández labs for assistance with data collection and cataloguing of specimens as well as helpful discussions surrounding the development of this manuscript. We would like to acknowledge Kati LaSance and the Vontz Center Imaging Laboratory at the University of Cincinnati for facilitating microCT scanning. We are thankful to Miguel García, the Nahá Lacandon community and Comisión Nacional de Áreas Naturales Protegidas (CONANP), for their help with our sampling collection. Finally, we acknowledge funding sources from Programa de Apoyo a Proyectos de Investigación e Innovación Tecnológica (IN212419) to CPO-G, Short stays program from AMC-FUMEC 2015 to CPOG, the National Institute of Health (NIDCR R01-DE025033) and National Science Foundation (DEB-1457630) to JBG, and ECOSUR (FID-784) to RR-H.

