## Supplementary figures and images for "A geographical cline in craniofacial morphology across populations of Mesoamerican lake-dwelling fishes"

### Supplemental Figure 1

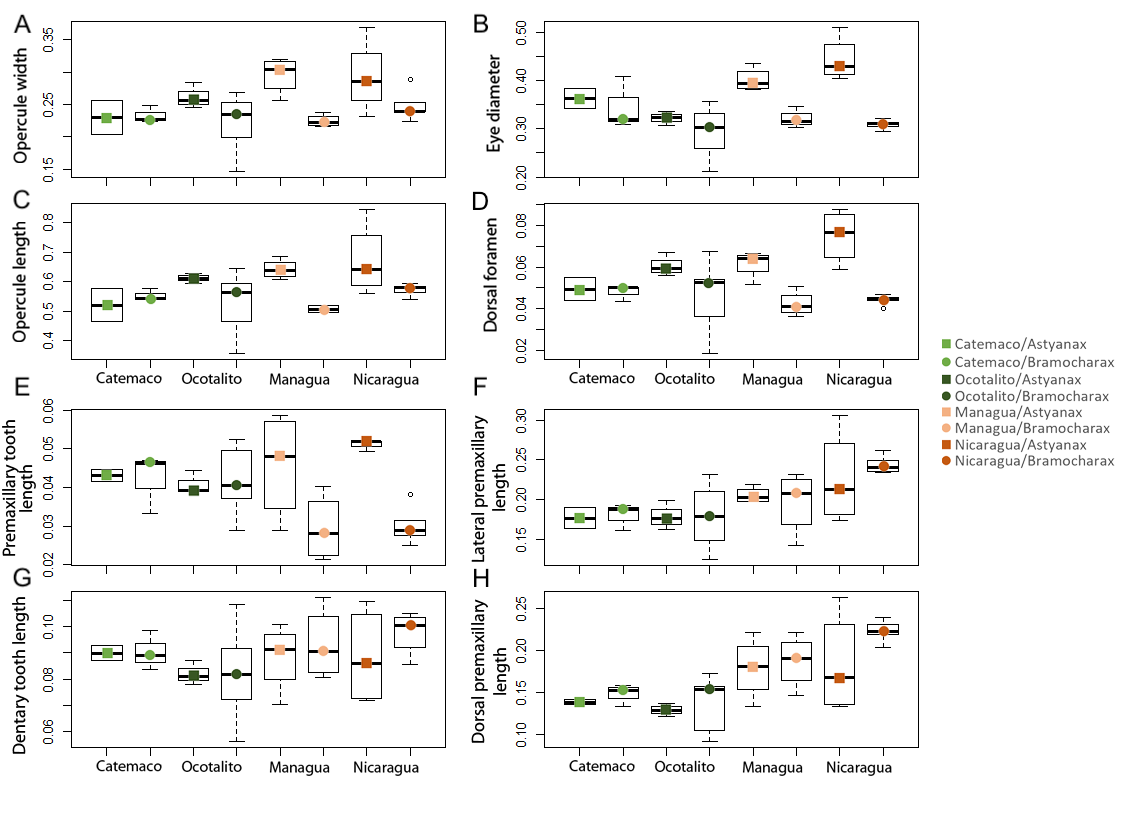

### Supplemental Figure 2

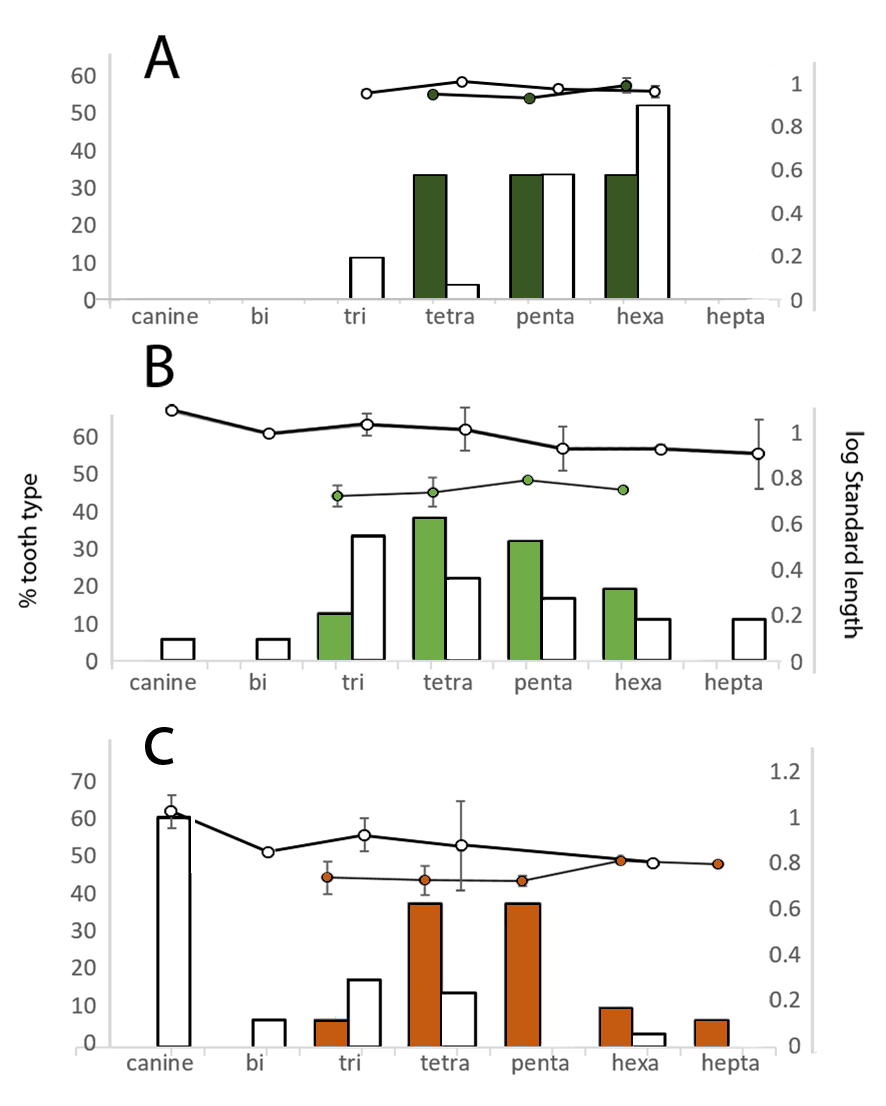
